# Acidosis-triggered fatty acid overload induces endothelial cell dysfunction

**DOI:** 10.64898/2026.07.09.737452

**Authors:** Sultan Al-Siyabi, Sébastien Ibanez, Kristian Serafimov, Justine Lallement, Dorothée Marchand, Florine Laloux, Céline Guilbaud, Delphine Demulder, Hanne Vlieghe, Saeid Moghassemi, Caroline Bouzin, Christiani Andrade Amorim, Olivier Feron, Chantal Dessy

## Abstract

Vascular ischemia is characterized not only by hypoxia but also by acidosis, which affects endothelial cells (ECs) due to increased H^+^ production from glycolysis and a deficit in H^+^ washout. We recently documented that an acidic environment facilitates the flip-flop transport of the non-ionized form of fatty acids (FAs) across the plasma membrane of cancer cells. In this study, we investigated how acidosis influences the capacity of highly glycolytic ECs to manage FAs and participates to endothelial dysfunction. We first tracked lipid droplet (LD) formation using Oil Red O staining and holotomographic microscopy. Purified monounsaturated oleate but also a mixture of FAs that reflect in vivo serum composition, resulted in dose- and time-dependent LD accumulation through FA transporter-independent mechanisms. Acid-exposed ECs exhibited enhanced mitochondrial respiration fueled by FAs, and endoplasmic reticulum (ER) stress, as indicated by the expression of ATF4 and CHOP. This phenotype was further associated with elevated reactive oxygen species production, which correlated with reduced nitric oxide (NO) availability. FA removal from EC culture media promoted lipolysis from LDs, supported by ATGL lipase induction which however slowed under acidic conditions. While ER stress persisted upon FA washout, NO availability was restored to levels comparable to those in FA-unexposed ECs. This observation coincided with dynamic mobilization of antioxidant defenses in acid-exposed ECs, as evidenced by low levels of reduced glutathione and enhanced cystine uptake, alongside a decrease in carnitine and FA-fueled mitochondrial respiration. Collectively, these data underscore the vulnerability of ECs to passive FA capture promoted by local acidosis, thereby contributing to a silent source of endothelial dysfunction in the postprandial state or during chronic exposure to elevated lipid levels.

## INTRODUCTION

Ischemia, characterized by insufficient blood flow to tissues, results in hypoxia, a condition that significantly restricts oxygen availability, thereby affecting the function of endothelial cells (ECs). ECs are inherently glycolytic and can readily adapt to reduced oxygen levels by predominantly relying on anaerobic metabolism, even under normal physiological conditions (1–3). This glycolytic dependence is thought to maximize oxygen delivery to the underlying parenchyma and to mitigate oxidative damage by maintaining reactive oxygen species (ROS) at manageable levels. An outcome of increased glycolytic flux in response to decreased pO_2_ is lactate production, accompanied by proton release. These protons accumulate locally due to the ischemic vasculature’s diminished capacity to clear them. Although a proton-enriched environment is frequently regarded as a collateral consequence of hypoxia, the decrease in extracellular pH (pHe) in ischemic tissues can be profound, with myocardial and cerebral interstitial pHe declining to approximately 6.0-6.5 during coronary occlusion or stroke (4, 5).

Multiple effects of ischemia on arterial tone and tissue perfusion have been linked to the acidic components of ischemia (6–13); however, definitive evidence distinguishing the effects of acidic pHe from the accompanying hypoxia or restricting them to the phase before reperfusion remains elusive. A notable exception pertains to the expression of several proton-sensing G protein-coupled receptors (GPCRs), particularly GPR4 and GPR68, which link changes in extracellular pH (pHe) to endoplasmic reticulum (ER) stress and a pro-inflammatory response (14–20). Proton-sensing GPCRs detect acidic pHe through alterations in the protonation and ionization of residues such as histidine, aspartate, and glutamate (21). These changes result in a reconfiguration of local electrostatics, hydrogen bonding, and associated conformational modifications (22).

By drawing an analogy with the protonation of protein residues, we recently demonstrated that fatty acids (FAs) can also benefit from the protonation of their carboxylate group in cancer cells (23, 24). Indeed, the resulting non-ionized form of FA is more likely to undergo the flip-flop mechanism, enabling their transport from the extracellular to the intracellular layer of the plasma membranes, where they are subsequently processed by FA-binding proteins in the cytosol. We identified this mechanism as a passive enhancement of lipid metabolism and peroxisomal oxidation in cancer cells located in acidic tumor regions (24). Here, we explored the potential for a similarly facilitated uptake of FA in acidotic endothelial cells (ECs) and its implications for their metabolism and nitric oxide (NO) production capacity.

It is important to emphasize that ECs have recently been identified as a barrier against lipotoxicity in the tissue parenchyma underlying blood vessels. Specifically, FA released from chylomicrons or lipoproteins by a lipase anchored to the luminal surface of the endothelium enter ECs and are incorporated as triglycerides into lipid droplets, thereby buffering the lipid surge associated with meals (25–28). Post-prandial lipolysis facilitated by adipose triglyceride lipase (ATGL) ensures the continuous and gradual release of free FAs, which can subsequently exit the endothelium to support the metabolism of parenchymal cells (29). Our findings indicate that extracellular acidosis leads to FA accumulation in ECs, which subsequently results in increased mitochondrial FA oxidation, endoplasmic reticulum (ER) stress, ROS production, and reduced nitric oxide (NO) availability. Furthermore, we demonstrated that the removal of the exogenous FA source restores NO production, although ER stress persists and fatty acid oxidation (FAO) is diminished. This identifies facilitated FA uptake as a significant contributor to endothelial dysfunction associated with acidosis.

## RESULTS

### ECs accumulate lipid droplets upon uptake of unsaturated fatty acids

We first aimed to establish an *in vitro* model of EC exposure to purified FA that mimics the *in vivo* conditions of postprandial exposure to mouse serum. We found that the serum of mice fed a high fat diet (HFD) resulted in the accumulation of lipid droplets (LDs), as evidenced by Oil Red O (ORO) staining, an effect that was further enhanced after 24h exposure (**Figures 1A-B**). The serum of mice on normal diet led to a less pronounced and delayed LD accumulation (**Figures 1A-B**). We next documented the capacity of monounsaturated fatty acid (MUFA) oleate to recapitulate these effects in a dose- and time-dependent manner (**Figures 1C-D**). The dose of 500 µM and 1000 µM were selected from a broader range of FA concentrations (not shown) to reflect the effects observed with the serum of mice on normal diet and HFD, respectively. We also observed that the addition of the polyunsaturated fatty acid (PUFA) docosahexaenoate (DHA) achieved approximately half the OA effects while under the same conditions the saturated fatty acid (SFA) palmitate failed to promote LD formation (**Figures 1E-F**), suggesting that the fate of SFA was primarily mitochondrial FAO.

**Figure 1.**
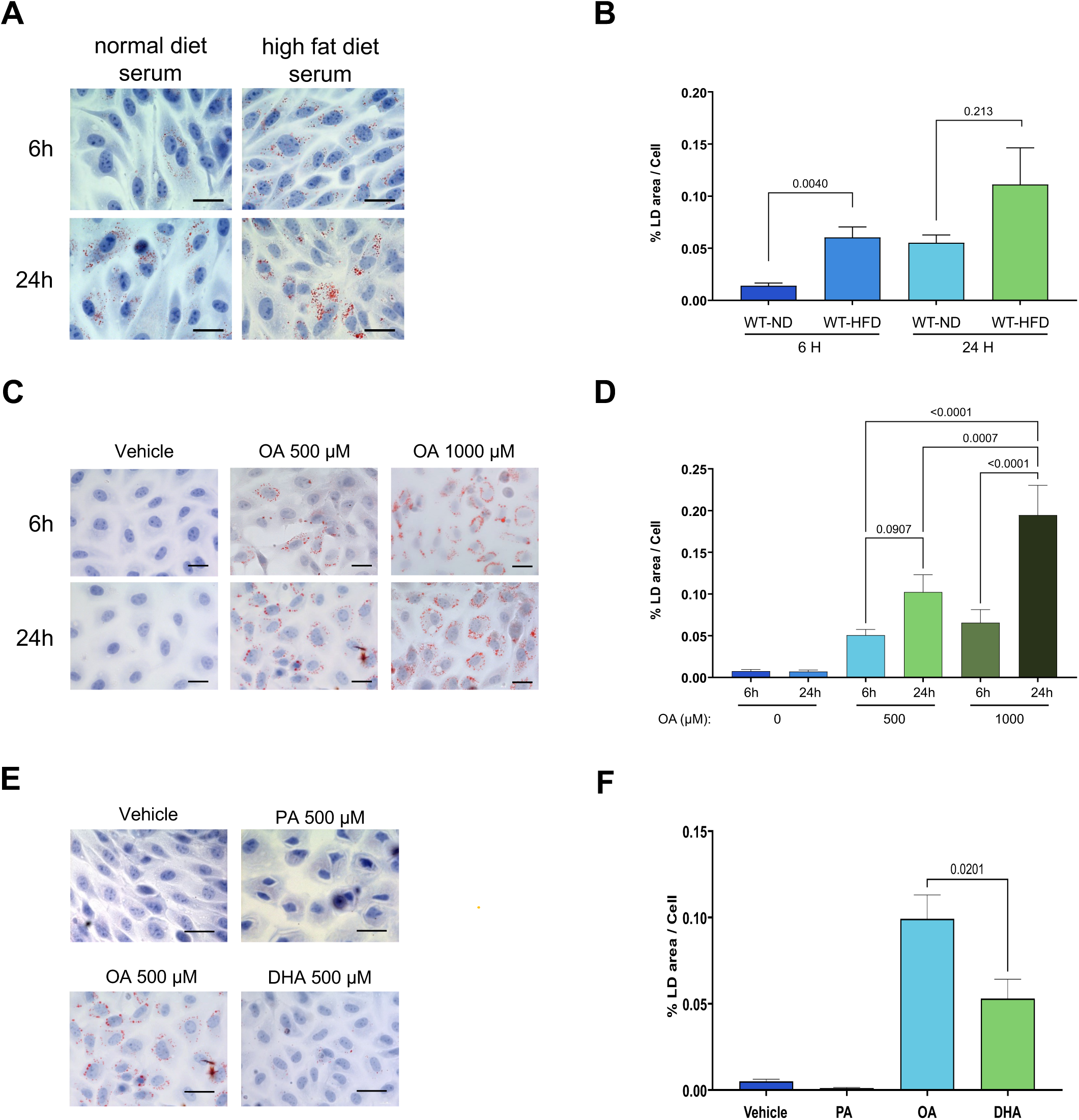
Lipid droplet accumulation in ECs exposed to FAs. Representative picture (A, C, E) and quantification (B, D, F) of lipid droplets (LDs), as evidenced by Oil Red O (ORO) staining in ECs. **(A-B)** LD accumulation in ECs exposed to serum from mice fed a high-fat diet (HFD) or normal diet (ND) for 24 hours (n=4). **(C-D)** Dose- and time-dependent effects of EC exposure to OA on LD formation; note that OA concentrations of 500 µM and 1000 µM lead to similar LD accumulation as ND and HFT mouse, respectively (n=5-9). **(E-F)** Comparison of LD formation in response to 24h exposure to 500 µM OA, palmitate (PA) and docosahexaenoate (DHA) (n=7).

### Extracellular acidosis enhanced FA uptake and LD accumulation in ECs

An extracellular pHe as observed in ischemic tissues has the potential to influence the ionization extent of the carboxylate group of fatty acids, and thereby to facilitate its transport across lipid bilayers. To investigate the impact of an acidic environment on the ability of ECs to manage FA overload, we cultured ECs at acidic pHe levels of 6.9 and 6.5, compared to a physiological pHe of 7.4. The accumulation of lipid droplets was significantly enhanced in correlation with the extent of acidosis (**Figures 2A-B**). We observed that this accumulation occurred rapidly, with lipid droplets readily detectable after 6 hours (**Suppl. Figures 1A-B**). Furthermore, to track this preferential FA uptake at earlier time, we used holotomographic microscopy for live tracking of LDs. This marker-free approach confirmed the pHe-dependent LD formation in ECs (**Figures 2C-D and Suppl. Movie 1**), and importantly that this was clearly apparent after 30-60 min precluding any transcriptional adaptation to acidic pHe (**Figure 2D**). We next examined whether FA transporters were involved in the observed increased FA uptake into ECs. Using SSO and FATP1-IN-1 to block CD36 and FATP1, respectively, we discovered that inhibition of FA transport significantly reduced the formation of LDs at pHe 7.4 whereas a substantial portion of LD formation at acidic pHe remained resistant to transport inhibition (**Figures 2E-F**). Finally, we explored the possibility of competition among different FA subtypes for uptake and subsequent storage. We used a physiologically relevant mixture of FAs including SFAs (3% C14:0, 25% C16:0 and 12% C18:0 for a total of 40%), MUFAs (4% C16:1, 41% C18:1, for a total of 45%) and PUFAs (12% C18:2, 3% C18:3 for a total of 15%). This more balanced mixture of FAs confirmed the pHe-dependent accumulation of LDs in ECs (**Figures 2G-H and Suppl. Figures 1C-D**). Collectively, these data support an acid-driven passive diffusion of FAs, likely resulting from the neutralization of their carboxylate groups at acidic pHe, as previously documented in cancer cells (24).

**Figure 2.**
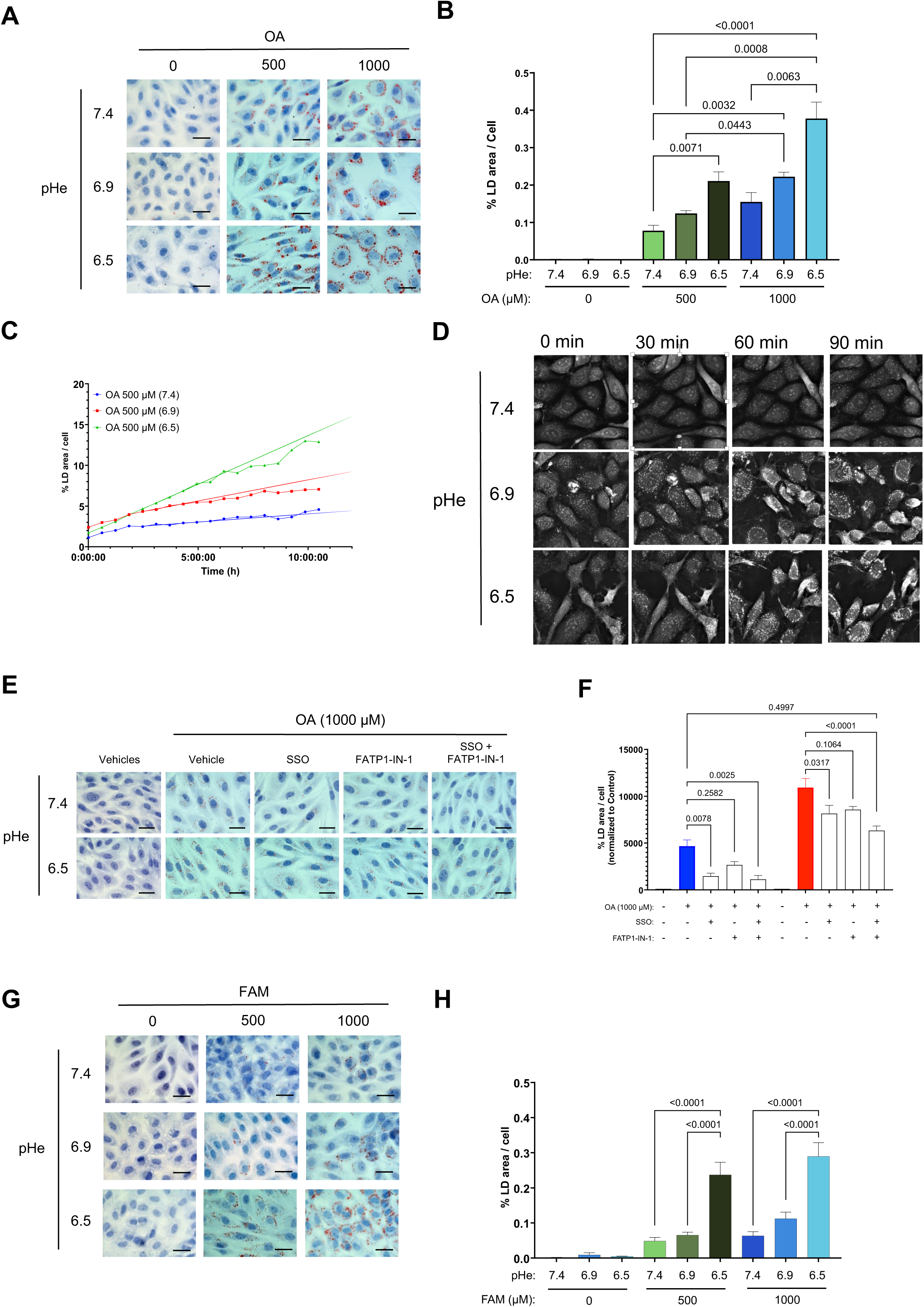
Acidic pHe enhances FA uptake and lipid droplet accumulation in ECs. **(A-B)** Representative ORO staining pictures and quantification of LDs in ECs cultured for 24 hours in the presence of the indicated concentration of OA at acidic pHe levels of 6.9 and 6.5 *vs.* pHe of 7.4 (n=4). **(C-D)** Time course measurements and representative pictures of LD formation in response to 500 µM OA at the indicated pHe as determined by holotomographic microscopy (n=4). **(E-F)** Representative ORO staining pictures and quantification of LDs in ECs cultured for 24 hours in the presence of 1 mM OA at pHe 7.4 and 6.5 in the presence of FA transporter blockers SSO and FATP1-1-IN (n=4). **(G-H)** Representative ORO staining pictures and quantification of LDs in ECs cultured for 24 hours in the presence of the indicated concentration of a FA mixture at acidic pHe levels of 6.9 and 6.5 *vs.* pHe of 7.4 (n=4).

### Extracellular acidosis and FA overloading heighten ER stress and ROS production in ECs

We next aimed to evaluate the extent of stress experienced by ECs in response to the enhanced FA uptake at acidic pHe. We investigated changes in the expression of ATF4 and CHOP, critical markers of endoplasmic reticulum (ER) stress (i.e., the site where LDs are formed) after 6h and 24h FA exposure (**Figures 3A-B**). We observed a time-dependent increase in ATF4 in ECs maintained at acidic pHe, even in the absence of FA addition, indicating that acidosis *per se* is a source of ER stress in ECs (**Figure 3C**, 6h- (left) and 24h-treatment (right)). CHOP levels similarly increased at acidic pHe but more prominently when OA or FAM was added (**Figure 3D**, 6h- (left) and 24h-treatment (right)). Notably, oleate exposure led to a more pronounced and lasting CHOP induction than that observed with the FA mixture (**Figure 3D**, right panel). We then examined whether this ER stress response could be associated with ROS production as measured by DCFA-based flow cytometry. We found a significant increase in ROS production in ECs maintained at acidic pHe compared to cells maintained at pHe 7.4 (**Figure 3E**). To further mimic a situation where ECs are subjected to chronic exposure to oxidative stress, we repeated these experiments in the presence of an external source of H_2_O_2_. Under these conditions, we observed that acid-exposed, FA-loaded ECs exhibited markedly elevated levels of ROS, indicating that their antioxidant defenses were less effective at mitigating the accumulation of oxidative stress (**Figure 3F**).

**Figure 3.**
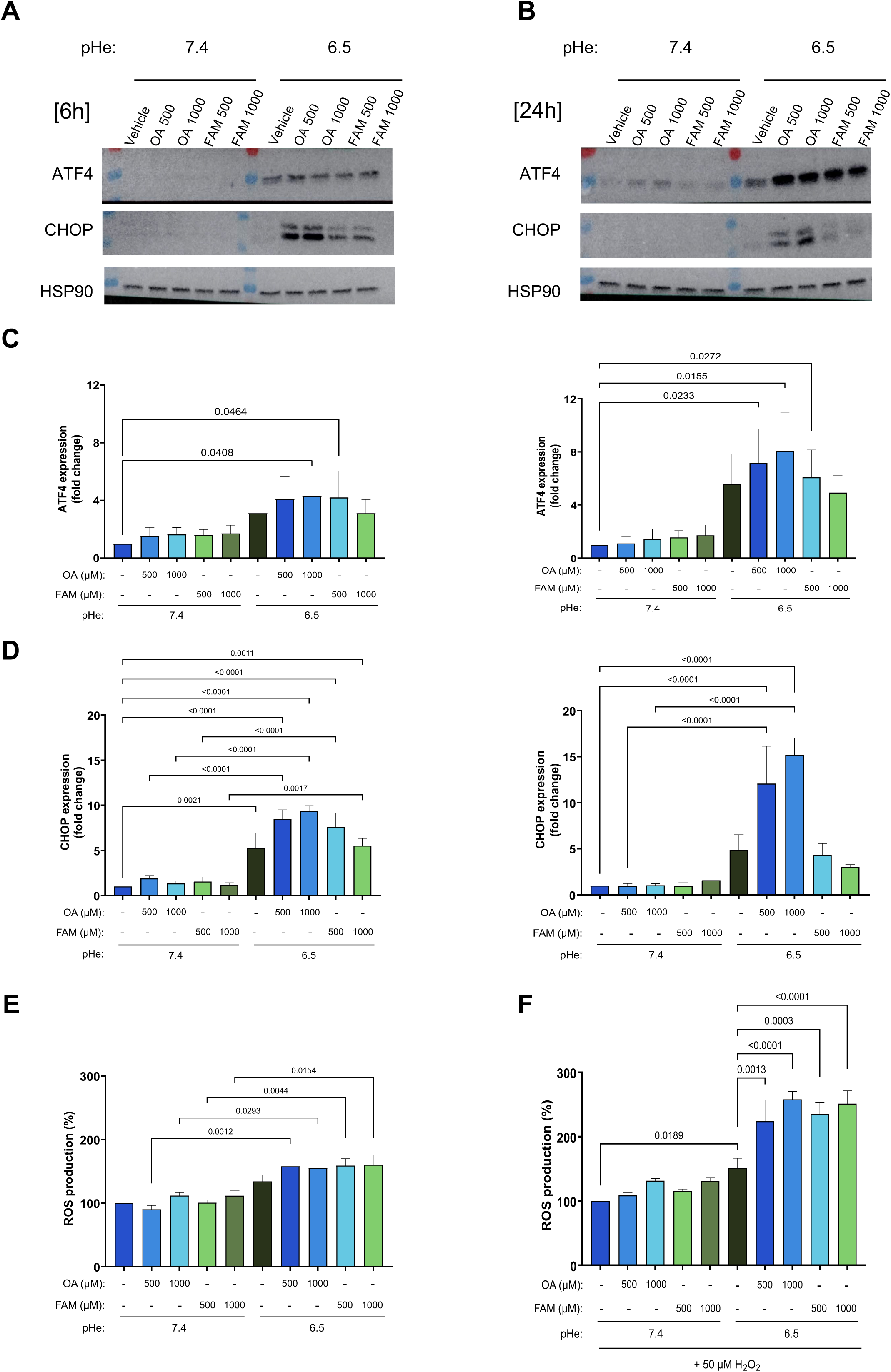
Acidosis-stimulated ER stress and ROS production in ECs exposed to FAs. Representative immunoblots and quantification of ATF4 and CHOP signals in ECs cultured in the presence of the indicated concentrations of OA or FA mixture at pHe 7.4 and 6.5. **(A-B)** ATF4 and CHOP immunoblots after 6 hours (A) and 24 hours (B) exposure to FA. **(C)** ATF4 expression levels after 6h (left) and 24h (right) (n=4). **(D)** CHOP expression levels after 6h (left) and 24h (right) (n=4). **(E)** ROS production in ECs cultured for 24h in the presence of the indicated concentrations of OA or FA mixture at pHe 7.4 and 6.5, either in basal conditions (left) or in the presence of H_2_O_2_ (right) (n=4).

### Extracellular acidosis and FA overloading reduce NO production and promote mitochondrial respiration in ECs

To verify whether the enhanced ROS production could lead to a reduced NO availability, we used EPR to directly measure NO in the extracellular medium. We first identified a significant reduction in NO release from ECs maintained at acidic pHe (vs pHe 7.4) (**Figure 4A**) whereas eNOS expression was not altered, as revealed by immunoblotting (**Suppl. Figures 2A-B**). Medium supplementation with OA or FAM induced a significant reduction in NO availability in ECs, regardless of the pHe (**Figures 4B-C**). We then reasoned that the combined effects of reduced availability of NO, a known inhibitor of respiratory chain complexed (30), along with enhanced FA fueling for oxidation, could result in increased mitochondrial activity. Using Seahorse technology, we confirmed that EC exposure to OA at pHe 6.5 resulted in a highly significant increase in both basal and maximal mitochondrial respiration (**Figures 4D and 4F-G**). To assess whether the increased mitochondrial activity was a result of FAO, we repeated these experiments in the presence of etomoxir, a CPT1 inhibitor (**Figure 4E**). We found that both basal and maximal OCR were significantly inhibited by etomoxir under both pHe conditions (**Figures 4E-G**). Of note, maximal respiration was elevated in ECs maintained at pHe 6.5 regardless of the presence of FA (**Figure 4G**), supporting an acidosis-induced metabolic shift toward oxidative phosphorylation (OXPHOS).

**Figure 4.**
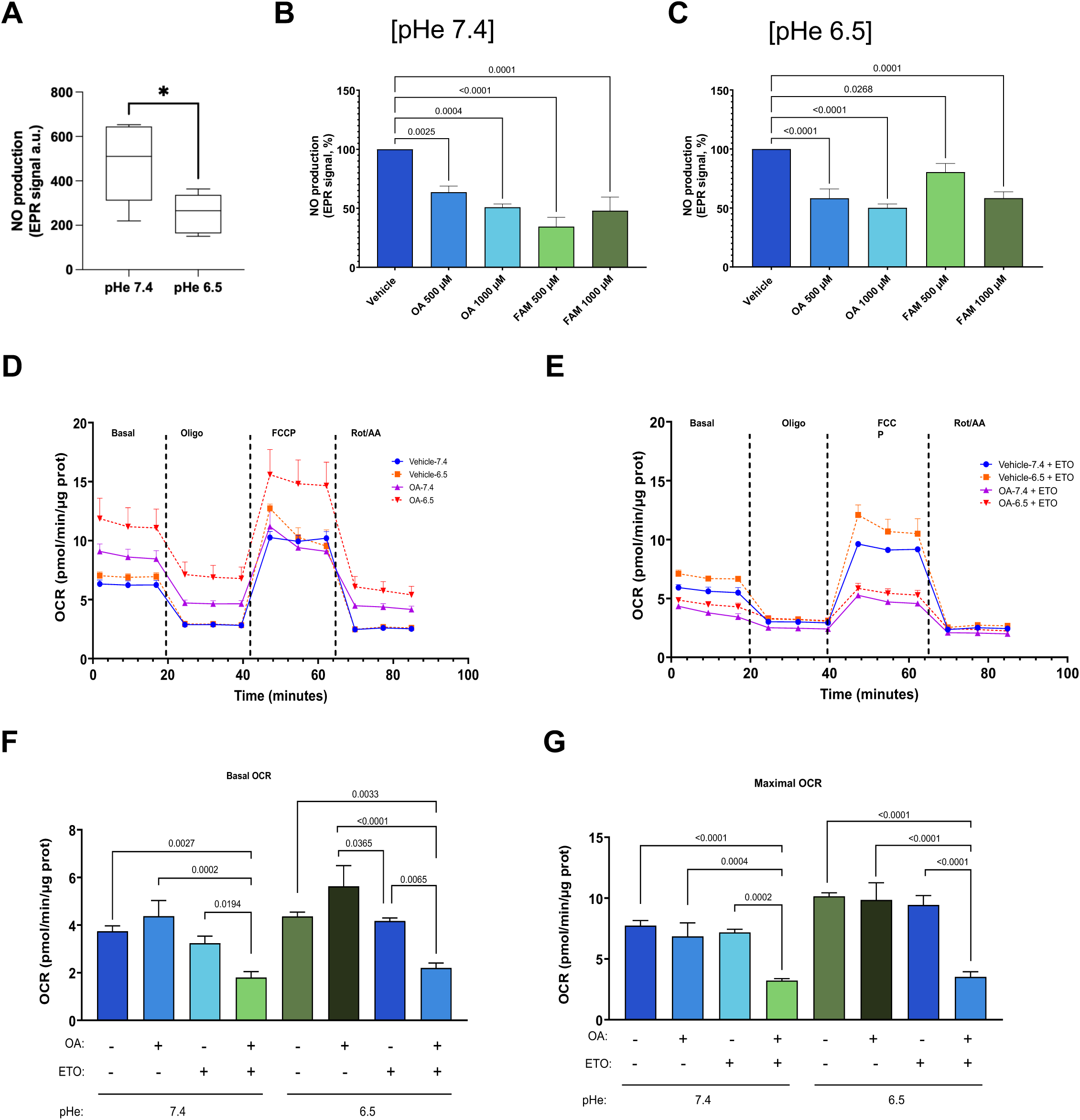
FA exposure accentuates acidosis-induced reduction in NO availability and increase in mitochondrial respiration in ECs. **(A)** EPR-based measurements of NO availability in ECs exposed at pHe7.4 and 6.5 (n=8). **(B-C)** Effects of 6h exposure to the indicated FA concentrations at pHe 7.4 (B) and pHe 6.5 (C), expressed as % of the NO availability determined in the absence of FA at pHe 7.4 and 6.5, respectively (n=4). **(D-E)** Seahorse measurements of O_2_ consumption rate (OCR) in ECs exposed to 500 µM OA (or vehicle as controls) at pHe 7.4 and 6.5, in basal conditions (D) or in the presence of CPT1 inhibitor etomoxir (E). Bar graphs depicting the basal **(F)** or maximal **(G)** OCR values in the experimental conditions described in D-E (n=6).

### ER stress persists in acid-exposed ECs throughout a slowed release of stored FA

In the next series of experiments, we examined the reversibility of the pathological EC phenotype induced by FA overload under acidic conditions. We found that FA wash-out stimulated lipolysis as evidenced by the time-dependent reduction in LD density (**Figures 5A-B**). Notably, the decrease in LD content was more rapid in ECs pre-challenged with FA at physiological pHe (**Figure 5A**) compared to those at acidic pHe (**Figure 5B**), and slightly more efficient from acid-exposed ECs exposed to FAM (*vs.* OA) (**Figure 5B**). In parallel, we documented the upregulation of the lipase ATGL in acid-exposed ECs as early as 6 hours after FA wash-out (**Suppl. Figures 3A-B**), which continued after 24 hours in ECs pre-challenged by FAM but not OA (**Figures 5C-D**). Interestingly, we observed that the induction of ATF4 and CHOP was detectable in FA pre-challenged acid-exposed ECs (*vs.* ECs maintained at pHe 7.4) 6 and 24 hours post-washout (**Suppl. Figures 3C-E** and **Figures 5E-G**, respectively); elevated CHOP expression persisted after 24 hours to a greater extent in acid-exposed ECs pre-loaded with OA (*vs.* FAM) (**Figures 5E and 5G**), aligning with the lesser ATGL induction in this condition (see Fig. 5C).

**Figure 5.**
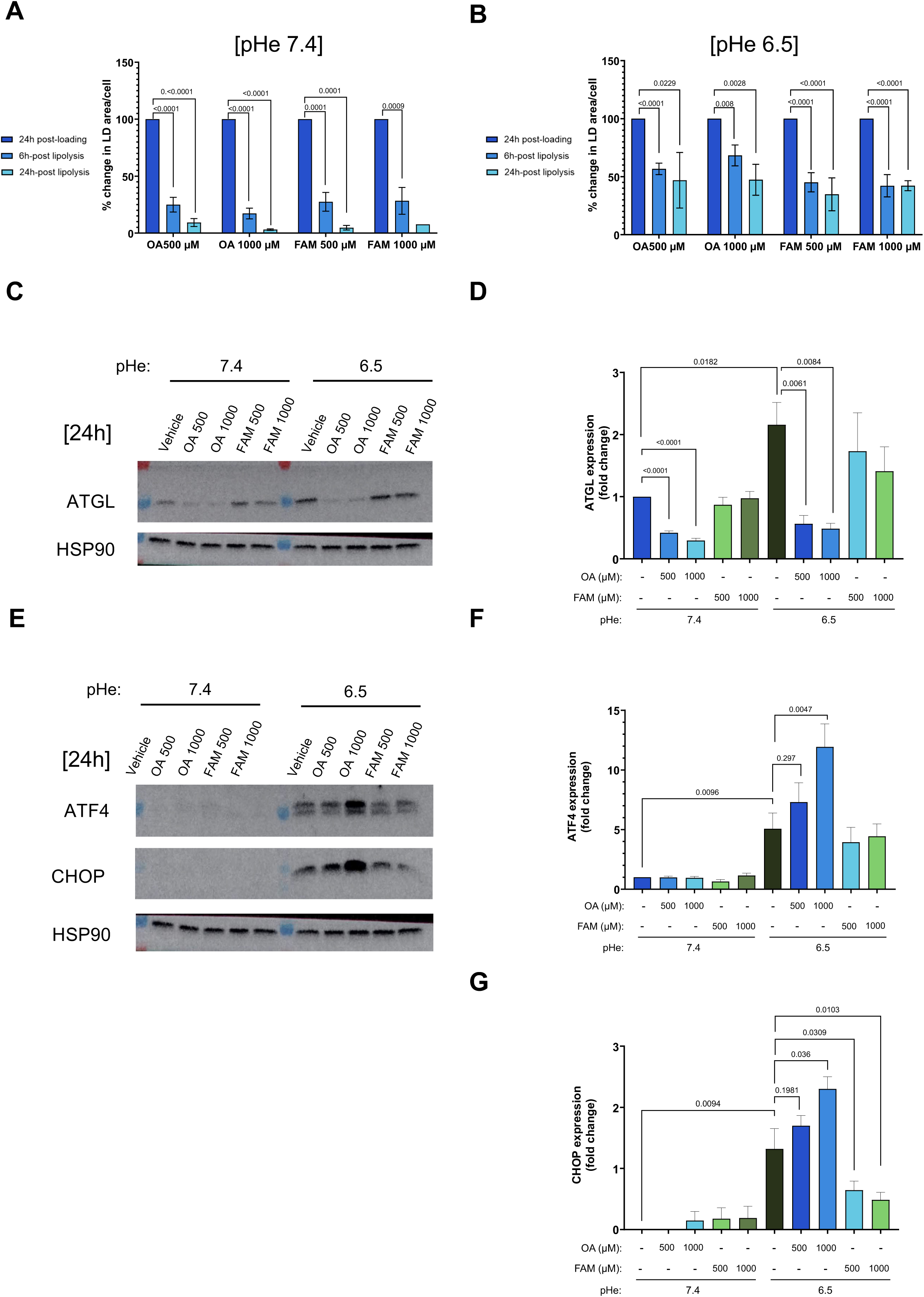
Effects of FA washout on the accumulation of lipid droplets and ER stress in ECs. **(A-B)** Changes in lipid droplet accumulation 6h and 24h following FA washout in ECs pre-challenged with the indicated FA for 24h at pHe 7.4 (A) and pHe 6.5 (B). **(C-D)** Representative immunoblots (C) and quantification (D) of ATGL signals following 24h FA washout in ECs pre-challenged with the indicated FA for 24h at pHe 7.4 and pHe 6.5. **(E-G)** Representative immunoblots (E) and quantification of ATF4 (F) and CHOP (G) signals in ECs following 24h FA washout in ECs pre-challenged with the indicated FA for 24h at pHe 7.4 and pHe 6.5 (n=4).

### Mobilization of antioxidant defenses and reduced respiration rate rescue NO availability during lipolysis

Finally, we evaluated the impact of FA washout on the capacity of ECs to produce NO. Surprisingly, while CHOP measurements had indicated that ECs were still facing stress despite wash-out (Figures 5E and 5G), we found that NO signal was not significantly altered anymore, regardless of the FA prechallenge (**Figures 6A-B**); eNOS immunoblotting confirmed that eNOS abundance remained constant following FA washout (**Suppl. Figures 6A-B**). To understand how the reversibility in NO availability was possible despite the ongoing the ER stress, we next examined the status of the antioxidant defenses. We found that GSH, the reduced form of glutathione, was lower at acid pHe following FA loading and remained at low level in the FA washout phase (*vs.* pHe 7.4) (**Figures 6C**). Levels of γ -glutamylcysteine, a precursor of GSH, were enhanced in the same conditions at pHe 7.4 indicating a progressive restoration of the anti-oxidant potential (**Figure 6D**). At pHe 6.5, γ -glutamylcysteine levels however remain low (**Figure 6D**), thereby confirming the persistence of the oxidative stress induced by acidic condition in the FA release stage. Interestingly, we also found a net increase in cystine after FA washout from acidic ECs (**Figure 6E**), suggesting a dynamic process in which cystine is utilized for the generation of GSH and immediate consumption to sustain an environment conductive to NO production.

**Figure 6.**
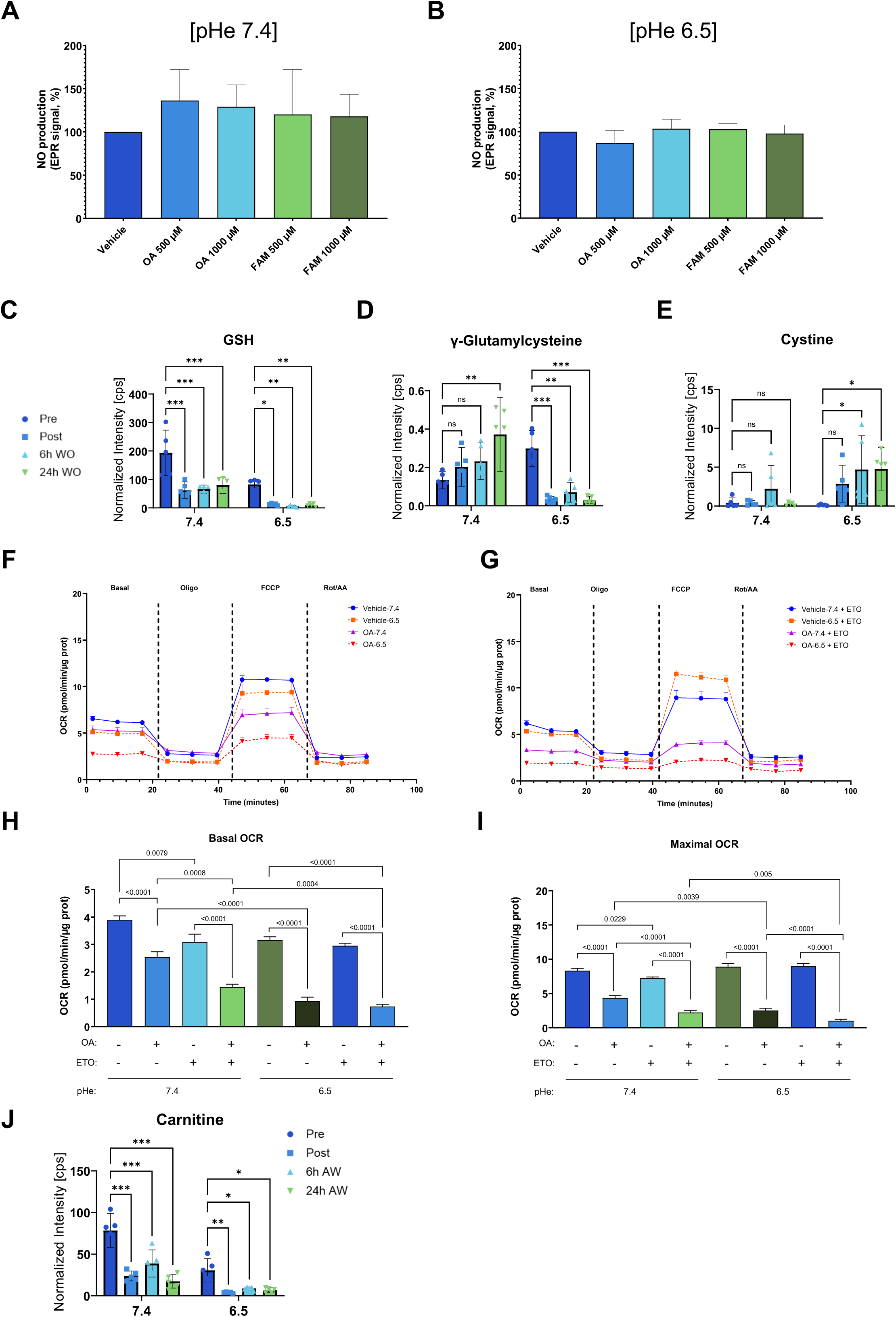
Effects of FA washout on NO availability and mitochondrial respiration in ECs. **(A-B)** Effects of 24h FA washout on NO availability as determined by EPR in ECs pre-challenged for 24h with the indicated FA concentrations at pHe 7.4 (A) and pHe 6.5 (B), data are expressed as % of the NO availability determined in the absence of FA at pHe 7.4 and 6.5, respectively (n=5). **(C-E)** LC-MS/MS quantification of reduced GSH (C), γ -glutamylcysteine (D) and cystine (E) in EC cultured at pHe 7.4 and pHe 6.5 before and after exposure to 500 µM OA, including following 6h and 24h FA washout (n=6). **(F-G)** Seahorse measurements of O_2_ consumption rate (OCR) in ECs 6h following FA washout in ECs pre-challenged with 500 µM OA (or vehicle as controls) at pHe 7.4 and 6.5, in basal conditions (F) or in the presence of CPT1 inhibitor (etomoxir) (G). Bar graphs depicting the basal **(H)** or maximal **(I)** OCR values in the experimental conditions described in F-G. **(J)** LC-MS/MS quantification of carnitine in EC cultured at pHe 7.4 and pHe 6.5 before and after exposure to 500 µM OA, including following 6h and 24h FA washout (n=6).

We finally investigated whether the restored NO production could inhibit OXPHOS, or if the continued release of free FA from LDs maintained elevated mitochondrial oxidative pathways. We found that both basal and maximal OXPHOS were reduced in ECs pre-exposed to OA, with a more pronounced decrease in ECs cultured at acidic pHe (**Figures 6F and 6H-I**); a slight inhibition of respiration by the CPT1 inhibitor etomoxir further indicated that FA supported residual OXPHOS (**Figures 6G and 6H-I**). This observation together with the slowed release of free FA from LD (see Figs 5A-B) led us to suspect a deficiency in the capacity of acyl-CoA to be taken up by mitochondria for catabolic oxidation. This was confirmed by a dramatic reduction in the levels of carnitine, which is essential for transporting acyl-CoA through mitochondrial CPT1, under acidic conditions (**Figure 6J**). This decreased availability of carnitine was observed following exposure of acid-exposed ECs to OA and persisted at very low levels at both 6 and 24h after FA washout (**Figure 6J**).

## DISCUSSION

The primary finding of this study is the demonstration of the role of extracellular acidosis, a hallmark of ischemic tissues, in promoting FA uptake into ECs, compelling them to adapt their metabolism but also impairing their ability to produce nitric oxide. Our work therefore adds the lipid nutrient layer to the understanding of how an elevated proton environment induces direct alterations in EC integrity and function (31). We identified a remarkable propensity of acidic pHe to accelerate FA uptake and initiate storage into lipid droplet within minutes, thereby simulating a postprandial state in an ischemic vessel. This enhanced passive acid-driven uptake appears largely independent of FA transporters, in contrast to the basal FA uptake process at neutral pHe. This study resonates with our previous work which documented that in tumors, local acidosis stimulates the uptake of the neutral form of FA, thereby facilitating the flip-flop transport of non-ionized FA across the plasma membrane (24). In cancer cells exposed to acidosis, we further showed that the consecutive FA overload results in a metabolic shift from glycolysis to enhanced peroxisomal FA oxidation (24). The situation in ECs differs, as these cells are primarily glycolytic and are transiently engaged in FA handling by taking up bloodborne FA, accumulating them as triglycerides and progressively releasing them into the subendothelial parenchyma. In response to acidosis, we demonstrated that ECs rapidly experience FA overload, exceeding their storage capacity in lipid droplets and fueling mitochondrial respiration (**Figure 7, middle panel**). The biological consequences of this abrupt phenotypic shift are immediate, evidenced by signs of ER stress, as indicated by the induction of ATF4 and CHOP, along with a significant increase in ROS production and reduction in NO production, as measured by EPR (**Figure 7, middel panel**).

**Figure 7.**
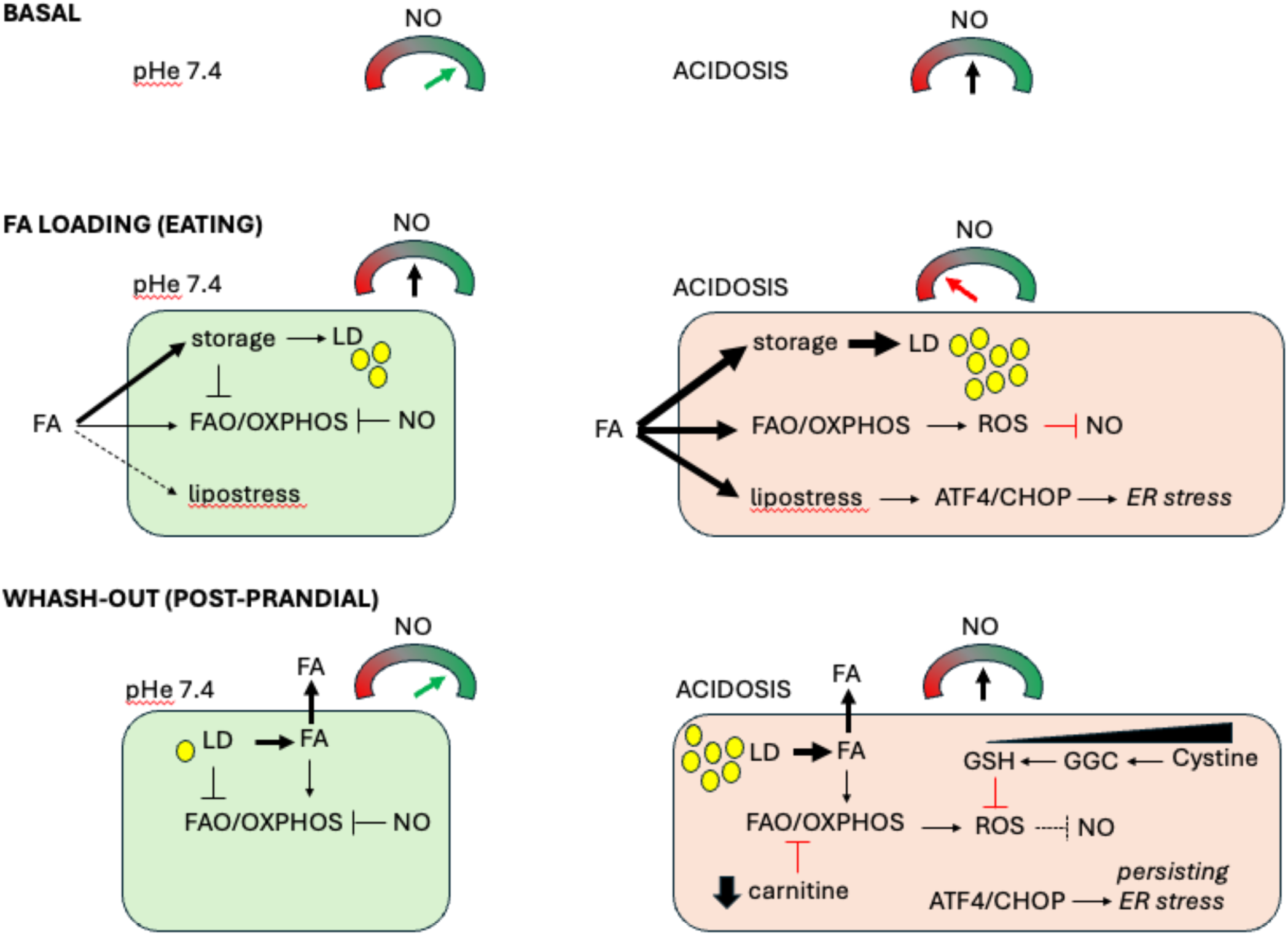

Our data uncover that both acidosis and FAs independently reduce the production of NO, yielding a significant aggregate decrease when both conditions coexist. Interestingly, upon removing the extracellular source of FA, we found that ECs restore their capacity to produce NO at levels comparable to those of ECs that were not pre-challenged by FA (even though it remains lower in absolute amounts under acidosis (Fig. 4A)). This observation can be analyzed in light of GSH-based antioxidant defenses. While reduced GSH and its precursor γ -glutamylcysteine levels remain near zero following FA washout, we observed a net increase in cystine. These data support a model in which GSH is readily consumed under these conditions to mitigate ROS production and thereby restore NO availability (**Figure 7, lower panel**). When examining the profiles of GSH, γ -glutamylcysteine and cystine, ECs exhibit a greater capacity to manage oxidative stress at physiological pHe, as evidenced by significantly elevated levels of GSH and increased levels of its precursor following FA washout. Notably, GSH and γ -glutamlycysteine levels are consistently higher than those observed at acidic pHe, regardless of the condition (i.e., before and after FA loading, and after FA wash-out). Another favorable context contributing to reduced ROS-related stress is the limited availability of carnitine, which persisted for at least 24 hours following FA washout, indicating that the transport of FA into the mitochondria was restricted, thereby mitigating ROS production (**Figure 7, lower panel**). This observation was substantiated by a net reduction in mitochondrial respiration, despite the continued presence of acidosis, and the associated plateau in lipid droplet lipolysis.

Our results regarding the restored nitric oxide (NO) production, which are at the limit of what the dynamic production of antioxidants can sustain, are based on a single instance of FA supplementation followed by a complete removal of the FA source. Therefore, repeated FA exposure, as naturally observed during postprandial periods, or chronic exposure to elevated FA levels may quickly exacerbate the situation and surpass the capacity of ECs to remain fully functional. This observation is further supported by our findings that FA pre-challenge in acid-exposed ECs induces ER stress, as indicated by the levels of ATF4 and CHOP, which are not resolved following FA washout (**Figure 7, lower panel**). Lipid droplets are known to play a central role in protecting ECs from intracellular free FA surges and lipotoxicity (29, 32, 33). However, when it comes time to release FA (following removal of extracellular FA), our data reveal that triglyceride stores are much higher at acidic pHe so that the amounts of free FA become rapidly toxic. Interestingly, we found that exposure to OA, but not to FAM, reduces the expression of ATGL, an observation that can be interpreted as an attempt to mitigate ER stress, which is actually more pronounced following OA exposure, as indicated by the larger CHOP signal compared to FAM conditions. Further research is needed to ascertain whether the excess OA, which is typically considered less detrimental than SFAs, disrupts the balance toward excessive membrane fluidity, thereby compromising EC integrity. Nevertheless, our findings indicate that by promoting FA overload, acidosis contributes to ER stress and an increased risk of cell death, adding to other mechanism previously identified, such as calcium overload (34). Beyond the immediate consequences for patients experiencing ischemia in the context of metabolic syndrome, obesity or diabetes (1, 32, 35, 36), the postprandial situation may pose a silent threat to individuals who otherwise appear to be in good health. Notably, humans spend more than half of the day in a postprandial state (37), and impaired vasoactivity has already been reported in young men during this period (38).

One limitation of our study is the use of macrovascular ECs instead of microvascular ECs. While the latter are more representative of the vascular bed involved in ischemic episodes, we chose macrovascular ECs due to their greater phenotypic stability in culture, including the maintenance of eNOS expression across passages. Still, the passive mechanism of stimulated FA uptake reported in our study is fundamentally governed by acid-base balance, making it applicable to any cell type subjected to acidosis, including microvascular ECs or myocytes. A second related limitation stems from the challenges of studying the relationship between acidosis and FA overload *in vivo* independently of other ischemia-associated events. In particular, the regional development of acidosis may not necessarily coincide with hypoxia. Local accumulation of H^+^ in ischemic zones can arise not only from hypoxia-driven glycolysis but also from a deficit in the washout of H^+^ resulting from respiration in the underlying parenchyma (i.e, CO_2_ + H_2_O → HCO_3_^-^ + H^+^). We previously reported such a lack of overlap between hypoxia and acidosis in tumor tissues (39).

Collectively, our data emphasize the significant role of FA overload in ECs exposed to acidosis as a critical contributor to endothelial dysfunction. This dysfunction arises not only from metabolic disturbances but also from the cellular stress responses induced by acidosis. Our findings reveal a new aspect of the broader impact of acidosis on vascular function besides the direct activation of proton-sensing GPCRs and ion channels, further exacerbating alterations in endothelial activity and integrity.

## METHODS

### Cell Culture

Bovine aortic endothelial cells (BAECs; Cell Applications, cat. no. B304-05) were cultured in T-75 flasks pre-coated with 0.2% (w/v) gelatin (Merck, cat. no. G1393) in endothelial cell growth medium (EGM; Cell Applications, cat. no. B211-500) at 37°C in a humidified incubator supplied with 5% CO₂. Cells were passaged at approximately 80% confluence, and experiments were conducted using cells between passages 3 and 8. Post-confluent BAECs were used throughout all experiments in this study. Mycoplasma contamination was routinely assessed using the MycoAlert Mycoplasma Detection Kit (Lonza, cat. no. LT07-318) according to the manufacturer’s instructions.

### Fatty Acid-BSA Complexes

Fatty acids (FAs) were complexed with fatty acid-free bovine serum albumin (BSA; Sigma-Aldrich, cat. no. A7030). The following FAs were purchased from Larodan (Solna, Sweden): oleic acid (OA; 18:1(n-9), cat. no. 10-1801-17), palmitic acid (PA; 16:0, cat. no. 10-1600-13), palmitoleic acid (POA; 16:1(n-7), cat. no. 10-1601-9), α-linolenic acid (ALA; 18:3(n-3), cat. no. 10-1803-9), linoleic acid (LA; 18:2(n-6), cat. no. 10-1802-13), stearic acid (SA; 18:0, cat. no. 10-1800-17), myristic acid (MA; 14:0, cat. no. 10-1400-13), and docosahexaenoic acid (DHA; 22:6, cat. no. 10-2206). FA stocks were prepared as described (40). Briefly, FAs were complexed with BSA at a 4:1 FA:BSA molar ratio, except for SA, which is prepared at a 2.5:1 ratio due to solubility limitations. Stock solutions were sterile-filtered (0.2 µm), aliquoted, stored at -80°C, and diluted into the experimental medium immediately before use.

### pHe-controlled media

Extracellular pH-controlled media were prepared by reconstituting DMEM powder (Merck, cat. no. D5030) in cell culture-grade ultrapure water supplemented with 5.5 mM glucose (Merck, cat. no. G8270), 2 mM GlutaMAX (ThermoFisher Scientific, cat. no. 35050038) and 0.4 mM phenol red (Merck, cat. no. P0290). To achieve a physiological pH of 7.4, 21.4 mM sodium bicarbonate (Merck, cat. no. S5761) was added. For acidic extracellular conditions, the basal medium was buffered with 25 mM HEPES (Merck, cat. no. H3375) and 25 mM PIPES sodium salt (Merck, cat. no. P2949). All complete media were supplemented with 10% fetal bovine serum (FBS, Merck, cat. no. F7524) and 1% penicillin-streptomycin (Gibco, cat. no. 15140-122). Target pH values (7.4, 6.9, and 6.5) were achieved by titration with NaOH or HCl and verified using a calibrated pH meter. All media were sterile-filtered (0.2 µm) under aseptic conditions, and pH stability was confirmed after a 24-h incubation at 37°C with 5% CO₂.

### Animal Studies and Plasma Collection

Five-week-old male C57BL/6J mice (n = 3–4/group) were fed either a standard diet (ND; SAFE A03, 13.5 kcal% fat) or a high-fat diet (HFD; D12492, 60 kcal% fat, Research Diets) for 24 weeks. Animals were housed in a temperature-controlled facility (12-h light/dark cycle) with ad libitum access to food and water. All procedures were approved by the Institutional Animal Care Committee of Université catholique de Louvain (UCLouvain; approval numbers: 2016/UCL/MD/003, 2020/UCL/MD/019, 2021/MD/02, and 2022/UCL/MD/62). Following the 24-week intervention, the mice were fasted for 12h with free access to water, then feeding them back to the original food. Three hours later, blood was collected at the postprandial stage. Mice were terminally anesthetized via intraperitoneal injection of ketamine/xylazine. Blood was collected via cardiac puncture into tubes containing 0.2% EDTA. Plasma was isolated by centrifugation (2800 rpm for 20 min at 4°C), snap-frozen in liquid nitrogen, and stored at −80°C until further analysis.

### EC Treatment with Fatty Acids

To model the *in vivo* postprandial FA surge, BAECs were treated for 6 or 24 h with 500 mM or 1000 mM of either individual FAs; oleic acid (OA), docosahexaenoic acid (DHA), or palmitic acid (PA), or a defined fatty acid mixture (FAM). The FAM was specifically formulated to reflect the most abundant lipid species in human serum, approximating circulating non-esterified fatty acid (NEFA) levels observed under normal and hyperlipidaemic conditions. The physiological FAM (total FA concentration: 500 µM) comprised: OA (41%), PA (25%), stearic acid (SA; 12%), linoleic acid (LA; 12%), palmitoleic acid (POA; 4%), myristic acid (MA; 3%), and alpha-linolenic acid (ALA; 3%). To simulate severe postprandial hyperlipidaemia, a corresponding pathological FAM was prepared using the same fatty acid composition ratios, but with a total FA concentration adjusted to 1000 µM. Where indicated, treatments were performed under controlled extracellular pH conditions (pH 7.4, 6.9, or 6.5).

### Oil Red O Staining and Quantification of Lipid Droplets

Bovine aortic endothelial cells (BAECs) were cultured on fibronectin-coated m-slides (ibidi, Gräfelfing, Germany). Following fatty acid treatment, the cells were washed once with phosphate-buffered saline (PBS) prior to fixation in 4% paraformaldehyde (PFA; Merck Millipore, Darmstadt, Germany) for 10 min at RT. Oil Red O (ORO) stock solution was prepared at 0.3% (w/v) (Merck, cat. no. O1391) in 100% isopropanol (VWR, Radnor, PA, USA, cat. no. 20922.411) and passed through a 0.2 µm filter. Immediately before use, an ORO working solution was prepared by mixing ORO stock solution with deionized water (3:1; v:v), followed by filtration through a 0.2 µm filter. Fixed cells were rinsed with double-distilled water and incubated in 60% isopropanol for 5 min. Following removal of isopropanol, cells were stained with ORO working solution for 15 min under gentle agitation. The cells were then thoroughly washed with double-distilled water, and nuclei were counterstained with hematoxylin (Dako, Glostrup, Denmark, cat. no. S3301) for 15 min. Slides were washed under running tap water to blue the hematoxylin stain and subsequently mounted using an aqueous mounting medium (Dako, cat. no. S3025). Bright-field images were acquired at 63X magnification using an Axio Imager Z1 microscope equipped with an ApoTome.1 system (Zeiss, Oberkochen, Germany). Lipid droplet area and nuclear count were quantified using ImageJ/Fiji software (NIH, Bethesda, MD, USA).

### Live-Cell Imaging of Lipid Droplet Formation via Holotomographic Microscopy

BAECs were seeded onto fibronectin-coated 35-mm glass-bottom imaging dishes (MatTek, Ashland, MA, USA, cat. no. P35G-1.5-14-C). For live-cell imaging, the culture medium was replaced with complete DMEM adjusted to a pH of 7.4, 6.9, or 6.5. The cells were treated with oleic acid (OA) at concentrations of 500 or 1000 µM. The dishes were then transferred to a top-stage microscope incubator (Nanolive, Ecublens, Switzerland) maintained at 37°C, 95% humidity, and 5% CO₂.

Live-cell imaging was performed using a 3D Cell Explorer holotomographic microscope (Nanolive) equipped with a 60× air objective . Holotomographic images were acquired over a 12-h period at 30-min intervals. Quantitative refractive index (RI) maps were reconstructed using the manufacturer’s built-in software. Lipid droplet accumulation was quantified utilizing the Smart Lipid Droplet Assay software module (Nanolive).

### Cell Viability

BAECs were seeded into gelatin-coated 96-well plates. The cells were treated with OA, PA, or FAM at concentrations of 500 and 1000 mM for 6 or 24 h in complete DMEM adjusted to pH 7.4, 6.9, or 6.5. Cell viability was evaluated using PrestoBlue Cell Viability Reagent (Thermo Fisher Scientific, Waltham, MA, USA, cat. no. A13262) according to the manufacturer’s protocol. Following a 1-h incubation at 37°C protected from light, fluorescence was measured at excitation/emission wavelengths of 560/590 nm using a SpectraMax i3 microplate reader (Molecular Devices, San Jose, CA, USA).

### Fatty acid uptake

BAECs were cultured on fibronectin-coated glass-bottom μ-slides (ibidi, Gräfelfing, Germany). Prior to fatty acid exposure, the cells were washed twice with complete Hanks’ Balanced Salt Solution (HBSS) and pre-incubated for 30 min at 37°C in complete HBSS containing one of the following: a CD36 inhibitor (15 µM sulfosuccinimidyl oleate sodium [SSO]; Merck, Darmstadt, Germany, cat. no. SML2148), a FATP1 inhibitor (2 µM FATP1-IN-1; MedChemExpress, Monmouth Junction, NJ, USA, cat. no. HY-141699), a combination of both inhibitors, or HBSS alone (vehicle control). Subsequently, the cells were exposed to 1000 µM oleic acid (OA) for 2 h at 37°C in complete DMEM, adjusted to pH 7.4 or 6.5, in the presence of the respective inhibitors. Following treatment, the cells were fixed and stained with ORO to visualize and quantify intracellular lipid accumulation.

### Nitric Oxide Bioavailability by EPR Spectroscopy

Nitric oxide (NO) production in BAECs was quantified by electron paramagnetic resonance (EPR) spectroscopy using Fe(II)-(DETC)₂ as a colloid spin trap, as previously described (41). Briefly, BAECs cultured in 100-mm dishes were treated with OA or FAM for 6 h. In a separate set of experiments, cells were treated with OA or FAM for 24 h, followed by incubation in lipolysis medium for another 24 h, as detailed below. Following treatment, the cells were washed twice with complete HBSS and incubated for 40 min at 37°C in Krebs-HEPES buffer containing 0.33% (w/v) BSA. Prior to the end of the treatment, Krebs-HEPES buffer (pH 7.4 or 6.5) and ultrapure HPLC-grade water were deoxygenated by bubbling with N₂ gas (30 min per 10–20 mL) and stored in a hypoxia chamber. The NO spin trap colloid (Fe(II)-(DETC)₂, 0.5 mM) was prepared immediately before use by dissolving diethyldithiocarbamic acid sodium salt (DETC; 0.451 mg/mL; Merck, cat. no. D3506) in deoxygenated Krebs-HEPES buffer and FeSO₄·7H₂O (0.278 mg/mL; Merck, cat. no. F7002) in deoxygenated water and subsequently mixing the two solutions gently. The spin trap colloid was then added to the cells (1:1 v/v), followed by incubation for 40 min at 37°C. Where indicated, NO production was stimulated using 2 µM calcium ionophore A23187 (Merck, cat. no. C7522).

Following incubation, the cells were scraped on ice into 100 µL of Krebs-HEPES buffer, snap-frozen in liquid nitrogen, and stored at -80°C until EPR measurement. For EPR measurements, samples were transferred to a liquid nitrogen finger Dewar (Noxygen, Elzach, Germany), and spectra were recorded on an X-band EPR spectrometer (Bruker Miniscope MS400; Magnettech, Berlin, Germany). The NO−Fe(DETC)₂ complex yields a characteristic three-line signal. NO production was quantified from the amplitude of the third hyperfine signal. Krebs-HEPES buffer (pH 7.4) contained 99.00 mM NaCl, 4.69 mM KCl, 1.03 mM KH₂PO₄, 1.20 mM MgSO₄, 25.00 mM NaHCO₃, 5.00 mM glucose, 20.00 mM HEPES, and 2.00 mM CaCl₂. For Krebs-HEPES at pH 6.5, NaHCO₃ was adjusted to 3.6 mM and NaCl to 120.4 mM to maintain osmolarity. All buffers were sterile filtered through a 0.22 mm membrane. Unless stated otherwise, chemicals were obtained from Sigma-Aldrich (St. Louis, MO, USA).

### Western Blotting

Following OA or FAM treatment, the cells were washed twice with ice-cold PBS supplemented with 1 mM sodium orthovanadate and lysed on ice in cold lysis buffer (Cell Signaling Technology, Danvers, MA, USA, cat. no. 9803) supplemented with protease inhibitor (Sigma-Aldrich, St. Louis, MO, USA, cat. no. P8340) and phosphatase inhibitor (Roche, Basel, Switzerland, Cat. no. 04-906-837-001). The lysates were clarified by centrifugation (10,000 × *g* for 10 min at 4°C).

Protein concentration was determined using a BCA protein assay (Pierce, Thermo Fisher Scientific, Waltham, MA, USA; cat. no. 23225). The samples were denatured in Laemmli buffer at 70°C for 5 min, and 20 µg of protein per sample was resolved by SDS-PAGE and transferred to 0.45 µm nitrocellulose membranes (Amersham, Cytiva, Marlborough, MA, USA; cat. no. 10600003). The membranes were blocked for 1 h in 5% non-fat milk in Tris-buffered saline with 0.1% Tween-20 (TBST) and incubated overnight at 4°C with primary antibodies diluted in 2.5% BSA in TBST.

After three 10-min washes with TBST, the membranes were incubated for 1 h at room temperature with horseradish peroxidase (HRP)-conjugated secondary antibodies (Jackson ImmunoResearch, West Grove, PA, USA; 1:10,000 in 5% milk/TBST). Following three TBST washes, signals were developed using an enhanced chemiluminescence (ECL) substrate (Amersham, cat. no. RPN2134) and captured using an Amersham Imager 600 instrument. Membranes were stripped as needed for reprobing. Molecular weights were estimated using a pre-stained protein marker (PageRuler; Thermo Fisher Scientific, cat. no. no. 26616). Band densitometry was performed using ImageJ/Fiji software (NIH, Bethesda, MD, USA), and phosphorylation levels were expressed as the ratio of phosphorylated protein to total protein, normalised to the BSA vehicle control (arbitrarily set to 100%). HSP90 was used as the loading control.

The following primary antibodies were used for immunoblotting: anti-HSP90 (1/10000, cat. no. 610419) and anti-eNOS (1/5000, cat. no. 610297) from BD Biosciences; anti-ATGL (1/2000, cat. no. 2138), anti-CHOP (1/2000, cat. no. 2895) and anti-ATF4 (1/2000, cat. no. 11815) from Cell Signaling Technology;

### Reactive Oxygen Species Quantification by Flow Cytometry

Intracellular reactive oxygen species (ROS) levels were quantified using flow cytometry with the fluorescent probe CM-H₂DCFDA. BAECs grown in 12-well plates were treated with OA, FAM, or FA-free BSA (vehicle control) for 24 h. For intracellular ROS detection, the cells were washed with complete HBSS and loaded with 30 µM CM-H₂DCFDA (Thermo Fisher Scientific, Waltham, MA, USA, cat. no. C6827) in HBSS for 25 min at 37°C in the dark. The cells were subsequently washed and incubated in phenol red-free complete DMEM for 15 min to allow probe de-esterification. The cells were then detached using 0.05% trypsin-EDTA, which was subsequently neutralized with 10% FBS in the HBSS. The cells were pelleted (1300 × g for 5 min), washed with HBSS, and resuspended in complete HBSS. Nuclei were stained with 0.5 mM DAPI (Merck, Darmstadt, Germany; cat. no. D9542). Fluorescence was measured using a BD FACSCalibur flow cytometer (BD Biosciences, San Jose, CA, USA). Both basal and stimulated (upon addition of 50 µM H_2_O_2_) ROS production were measured. Data were analyzed using FlowJo software (v. 10.10; BD Biosciences).

### Lipolysis Assays

Intracellular lipid droplet turnover and lipolysis were investigated using a modified protocol published by Luk et al. (42) Briefly, BAECs were treated with OA or FAM to induce intracellular lipid droplet accumulation. The cells were then briefly washed twice with complete HBSS and incubated in DMEM supplemented with lipid-depleted FBS (cat. no. S140L, Biowest).

### Measurement of Mitochondrial Respiration

Mitochondrial respiration was assessed by measuring the oxygen consumption rate (OCR) using a Seahorse XFe96 Analyzer (Agilent Technologies, Santa Clara, CA, USA). Briefly, ECs were seeded at a density of 20,000 cells/well in gelatin-coated 96-well Seahorse cell culture microplates. The cells were treated for 24 h in complete DMEM adjusted to pH 7.4 or 6.5 with 1 mM OA or FA-free BSA as vehicle control. In a separate set of experiments, ECs were treated for 24 h with 1 mM OA or FA-free BSA, after OA withdrawal, and incubated in lipolysis medium for another 6 and 24 h. Prior to the assay, the cells were equilibrated at 37°C in Seahorse XF DMEM assay medium supplemented with 1 mM sodium pyruvate, 2 mM L-glutamine, and 5.5 mM glucose (pH adjusted to 7.4 or 6.5). The Seahorse XF Cell Mito Stress Test cartridge was hydrated according to the supplier’s recommendations. OCR was recorded at baseline and after sequential injection of oligomycin (2.5 mM), FCCP (1.5 mM), and a mixture of rotenone and antimycin A (0.5 mM each). All experimental conditions were tested on the same plate using cells from a single passage in each experiment. The final OCR values were normalized to the total cellular protein content, which was determined post-assay using a BCA protein assay (Pierce, Thermo Fisher Scientific, Waltham, MA, USA, cat. no. 23225).

### Liquid Chromatography-Mass Spectrometry (LC-MS) Metabolomics

To investigate alterations of GSH, γ -glutamylcysteine and cystine, along with carnitine variations, ECs in response to OA treatment and subsequent lipid droplet mobilization under distinct extracellular pHe conditions, we performed LC-MS measurements according to the standardized protocol of UCL-MetlosLib (43). Briefly, ECs were seeded in 60-mm culture dishes and treated for 24 h with 1 mM OA at pHe of 7.4 or 6.5. The control cells were treated with FA-free BSA. To evaluate changes in the above parameters during lipolysis, a subset of OA pre-loaded cells for 24 h was subjected to OA withdrawal and incubated in the lipolysis medium for an additional 6 or 24 h.

### Statistical Analysis

All data are presented as the mean ± standard error of the mean (SEM). Statistical analyses were performed using GraphPad Prism software (version 10; GraphPad Software, San Diego, CA, USA). Comparisons among multiple groups were performed using a one-way or two-way analysis of variance (ANOVA) followed by an appropriate post hoc test for multiple comparisons. A P-value ≤ 0.05 was considered statistically significant. Sample sizes, defined as the number of independent biological replicates (n) are detailed in the figure legends.

## Acknowledgements

This work was supported by grants from the Fonds de la Recherche Scientifique (F.R.S.-FNRS, CDR J006722F and EOS O002522F) and an Action de Recherche Concertée ARC 23/28-132. SA-S is the recipient of a Sultan Qaboos University college of Medicine and Health Sciences fellowship. O.F. is a senior WELBIO investigator and C.D. is a F.R.S.-FNRS senior research associate.

