## Supplementary material for "Acidosis-triggered fatty acid overload induces endothelial cell dysfunction": Suppl. Figures

#### Suppl. Figure 1

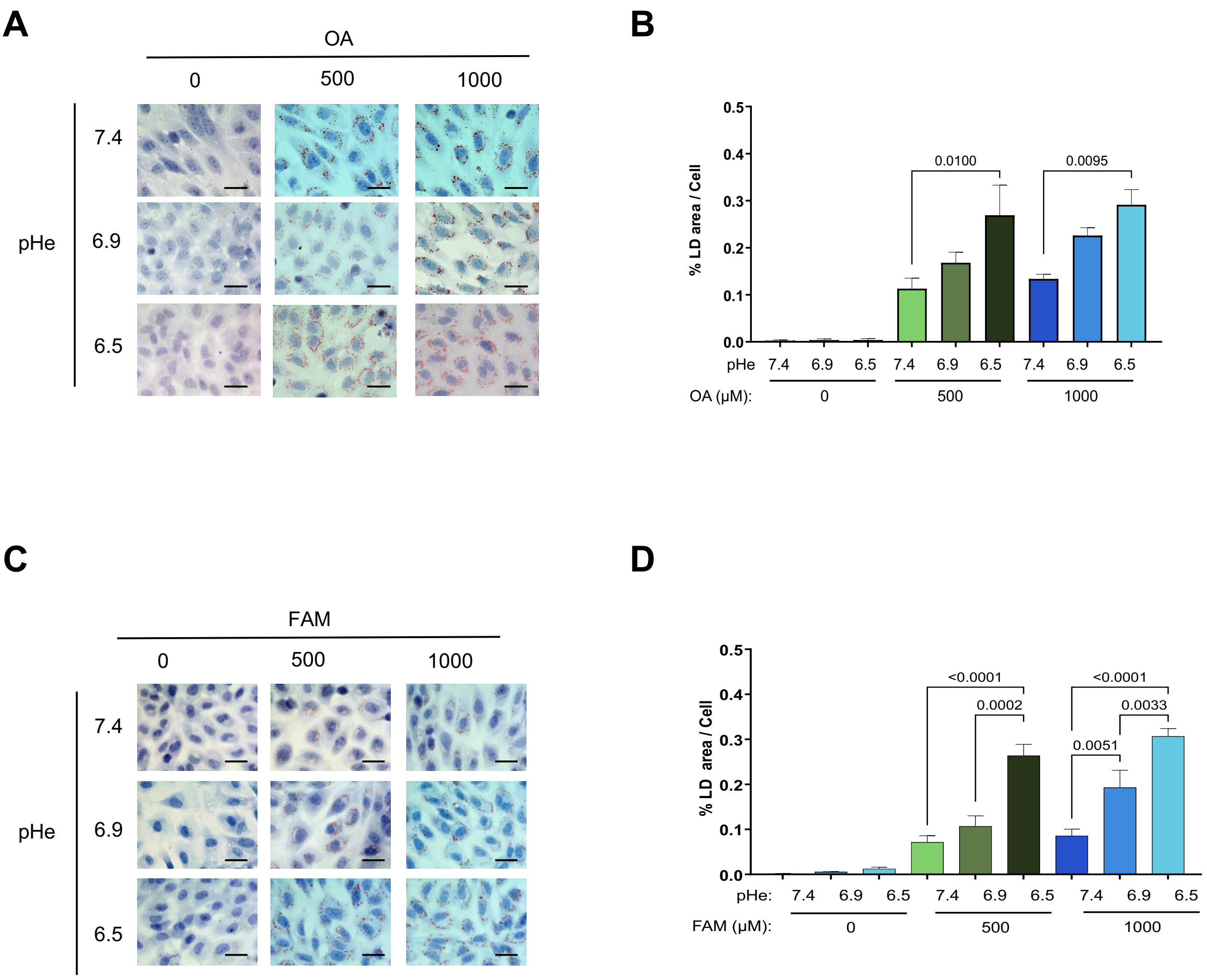


**Suppl. Figure 1. Acidic pHe enhances FA uptake and lipid droplet accumulation in ECs. (A-B)** Representative ORO staining pictures and quantification of LDs in ECs cultured for 6 hours in the presence of the indicated concentration of OA at acidic pHe levels of 6.9 and 6.5 *vs.* pHe of 7.4. **(C-D)** Representative ORO staining pictures and quantification of LDs in ECs cultured for 6 hours in the presence of the indicated concentration of a FA mixture at acidic pHe levels of 6.9 and 6.5 *vs.* pHe of 7.4 (n=4).

#### Suppl. Figure 2

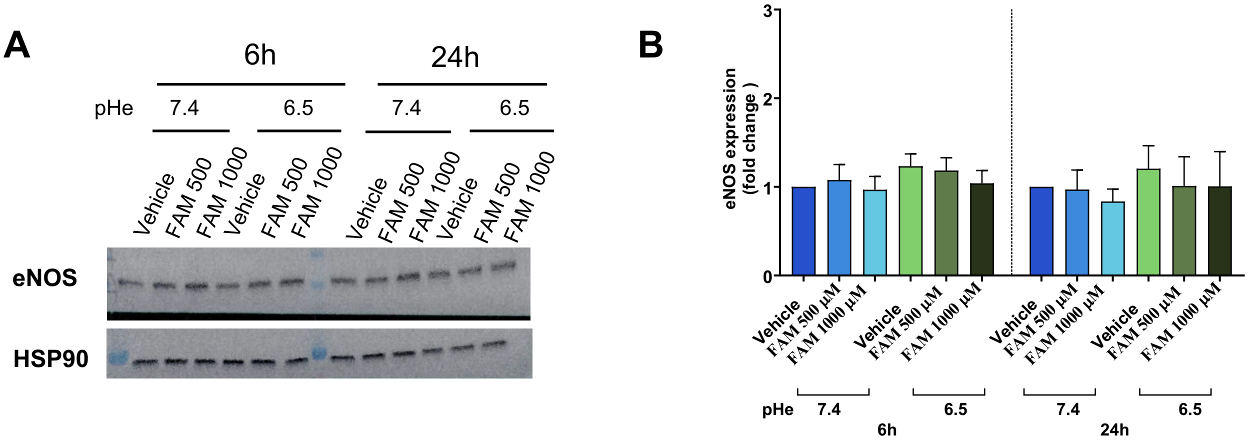


**Suppl. Figure 2. Acidic pHe and FA exposure do not impact on total eNOS abundance. (A-B)** Representative immunoblots (A) and quantification (B) of eNOS signals in ECs cultured for 6h and 24h in the presence of the indicated concentrations of FA mixture at pHe 7.4 and 6.5.

**Suppl. Figure 3**

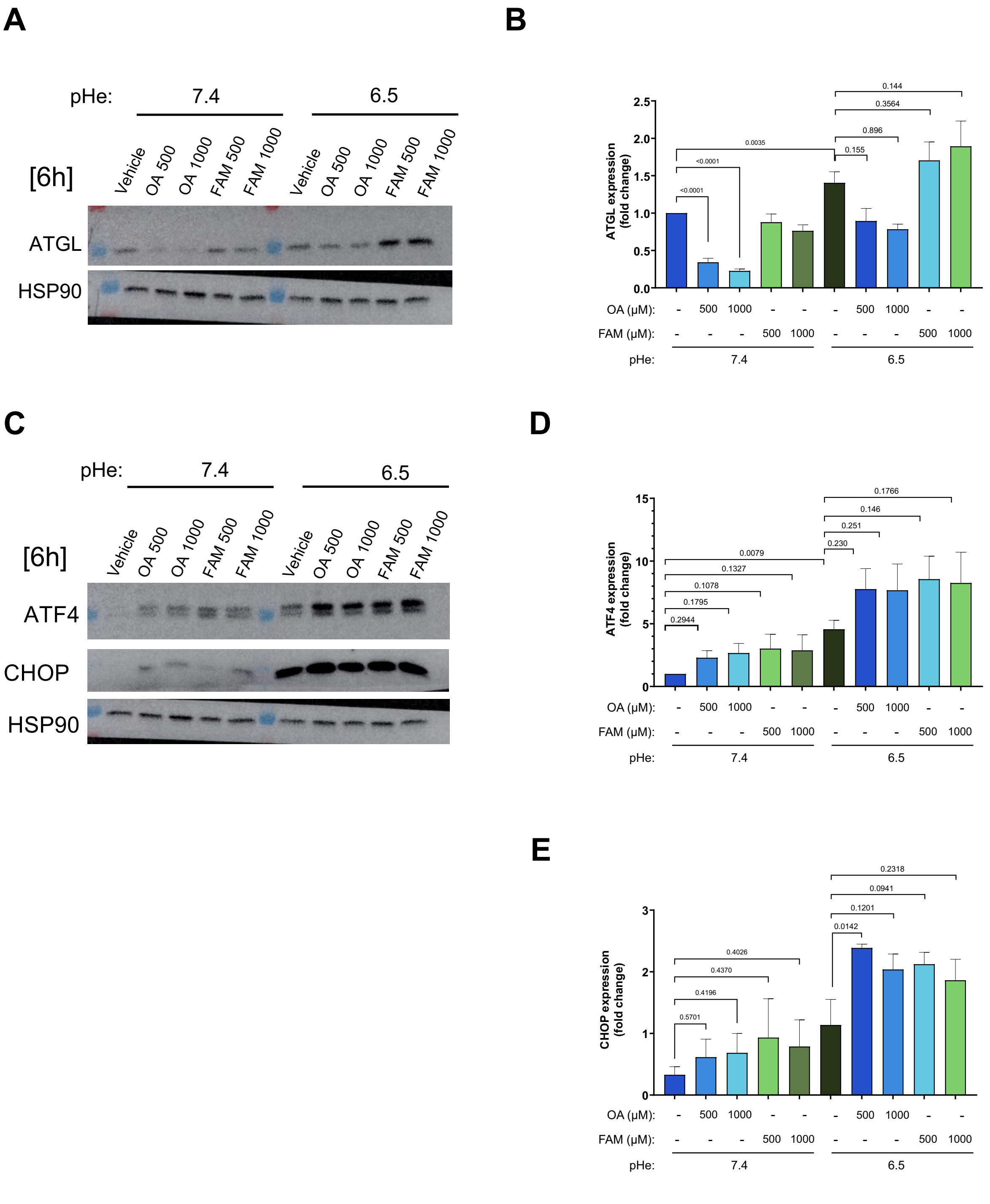


**Suppl. Figure 3. Effects of FA washout on the accumulation of lipid droplets and ER stress in ECs. (A-B)** Representative immunoblots (A) and quantification (B) of ATGL signals following 6h FA washout in ECs pre-challended with the indicated FA for 24h at pHe 7.4 and pHe 6.5. **(C-E)** Representative immunoblots (C) and quantification of ATF4 (D) and CHOP (E) signals in ECs following 6h FA washout in ECs pre-challended with the indicated FA for 24h at pHe 7.4 and pHe 6.5.

### Suppl. Figure 4

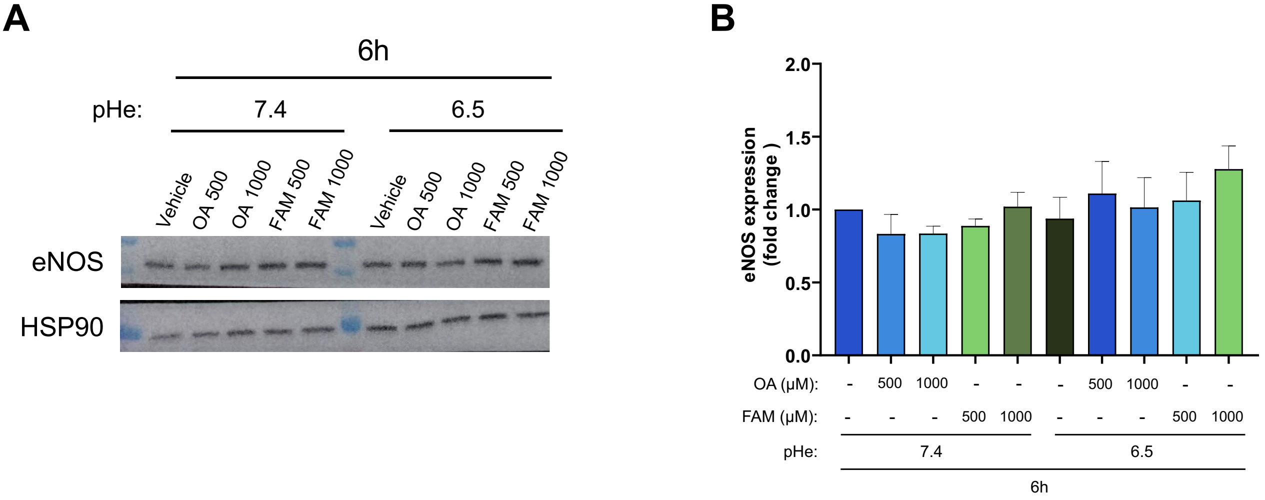


**Suppl. Figure 4. FA washout does not impact on total eNOS abundance. (A-B)** Representative immunoblots (A) and quantification (B) of eNOS signals in ECs following FA washout in ECs pre-challended for 6h and 24h in the presence of the indicated concentrations of OA or FA mixture at pHe 7.4 and 6.5 (n=4).
