## Supplementary material for "Acidosis-triggered fatty acid overload induces endothelial cell dysfunction": Suppl. movie 1

### Slide 1
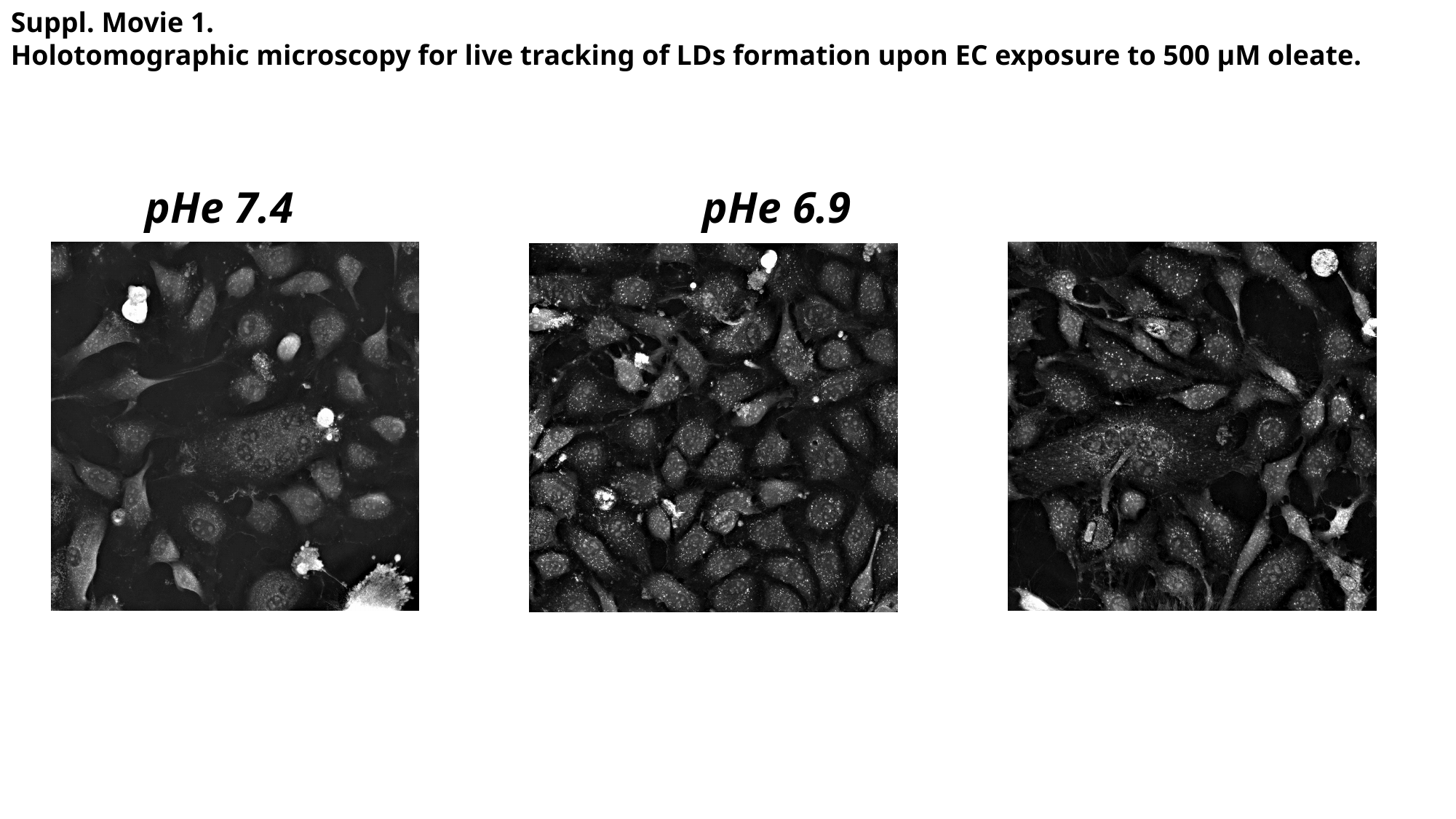

Suppl. Movie 1.
Holotomographic microscopy for live tracking of LDs formation upon EC exposure to 500 µM oleate.
pHe 7.4 pHe 6.9 		pHe 6.5
